# Visualization & Quality Control Tools for Large-scale Multiplex Tissue Analysis in TissUUmaps 3

**DOI:** 10.1101/2022.12.01.518716

**Authors:** Andrea Behanova, Christophe Avenel, Axel Andersson, Eduard Chelebian, Anna Klemm, Lina Wik, Arne Östman, Carolina Wählby

## Abstract

Large-scale multiplex tissue analysis aims to understand processes such as development and tumor formation by studying the occurrence and interaction of cells in local environments in e.g. tissue samples from patient cohorts. A typical procedure in the analysis is to delineate individual cells, classify them into cell types, and analyze their spatial relationships. All steps come with a number of challenges, and to address them and identify the bottlenecks of the analysis, it is necessary to include quality control tools in the analysis workflow. This makes it possible to optimize the steps and adjust settings in order to get better and more precise results. Additionally, the development of automated approaches for tissue analysis requires visual verification to reduce skepticism with regard to the accuracy of the results. Quality control tools could be used to build users’ trust in automated approaches. In this paper, we present three plugins for visualization and quality control in large-scale multiplex tissue analysis of microscopy images. The first plugin focuses on the quality of cell staining, the second one was made for interactive evaluation and comparison of different cell classification results, and the third one serves for reviewing interactions of different cell types.

## 1 Introduction

Understanding cell distributions and interactions in tissue plays a crucial role in order to fully comprehend organism development, healing, and homeostasis [1]. Particularly important for this is the spatial distribution of different cell types, which can be identified by various spatial omics techniques, such as multiplexed spatially resolved analysis of gene expression [2, 3, 4, 5] or multiplex immunohistochemical (IHC) staining microscopy [6, 7, 8, 9, 10]. Analysis of gene expression and IHC should be considered complementary techniques, where gene expression is related to the function of the cells and IHC approaches provide a direct understanding of marker protein expression, post-translational modifications, and sub-cellular localization [11]. It has also been shown that multiplex IHC has significantly higher performance than gene expression profiling for predicting objective response to a certain therapy [12]. Multiplex IHC makes it possible to identify multiple cell types in parallel and enables the study of cell-cell interactions and local cell environments. The first choice of analysis is often semi-automated visual/manual cell classification by intensity thresholding. There is a trend to move towards fully automated tools, not only to speed up analysis but also to reduce bias. However, the development of automated processing steps requires visual inspection and quality control tools to optimize settings and confirm the correctness. These tools help to increase the user’s trust in automated systems, especially if compared to some kind of validated ground truth.

More steps in an analysis approach bring more challenges, and there are several essential steps required in order to be able to quantify cell-cell interaction. The first step is IHC staining, coming with challenges including weak staining, high background intensity, over-staining or nonspecific staining [13, 14]. The second step, microscopy imaging, bears obstacles such as nonuniform illumination or low/high contrast [15]. Cell classification is the next step and brings challenges such as misclassification, and false negative classification, especially if cells are crowded and partially overlapping. The last step is the quantification of interaction, which can be done by various methods [16], and selecting which one to use might not be trivial. To conclude, there are many challenges in the analysis workflow and all these steps have a large impact on the final result. Careful quality control combined with visual assessment is necessary to compare and validate different options.

We present here three plugins for quality control and visual assessment of the analysis intermediate and final results. The first one is for quality control and comparison of cell staining. The second one is for quality control and comparison of cell classification. And the third one is for quality control and visualization of cell-cell interactions. We also provide a new approach to quantify cell-cell interactions that takes local tissue structures into account. Other tools for evaluating cell-cell interaction exist, such as ImaCytE [17], which can highlight the interaction of protein expression profiles in microenvironments. However, ImaCytE was developed for Imaging Mass Cytometry data, unlike TissUUmaps 3 [18] which is suitable for any type of marker data.

In this paper, present each of the plugins together with example data, and share the plugins via https://tissuumaps.github.io/.

## 2 Software and methods

In this section, we present newly developed plugins for the free and open-source software TissUUmaps 3 [18]. TissUUmaps 3 is a browser-based tool for fast visualization and exploration of millions of data points overlaying gigapixel-sized multi-layered images. TissUUmaps 3 can be used as a web service or locally on your computer and allows users to share regions of interest and local statistics. The three plugins are Visualization comparison and quality control of cell staining (StainV&QC), Visualization comparison and quality control of cell classification (ClassV&QC) and Visualization and quality control of cell-cell interactions (InteractionV&QC).

### 2.1 Overview

The typical steps from multiplex microscopy image data collection to quantitative analysis of cell-cell interactions are shown in Figure 1, together with the necessary quality controls and corresponding plugins. Initial steps, such as cell segmentation, feature extraction, and cell classification, require external tools such as CellProfiler [19, 20], CellProfiler Analyst [21] and QuPath [22]. Accumulation scores quantifying cell-cell interactions can be calculated by tools such as Squidpy [23] and histoCAT [24]. The results are visually inspected and checked for quality by the three plugins. Plugin *StainV&QC* compares raw data from different cores and the effect of image pre-processing methods. Plugin *ClassV&QC* visually verifies/discards classification results or compares classification results achieved by different manual and/or automated classification approaches. Plugin *InteractionV&QC* visually verifies/discards and investigates non-random tissue patterns and spatial cell-cell interactions.

**Figure 1:**
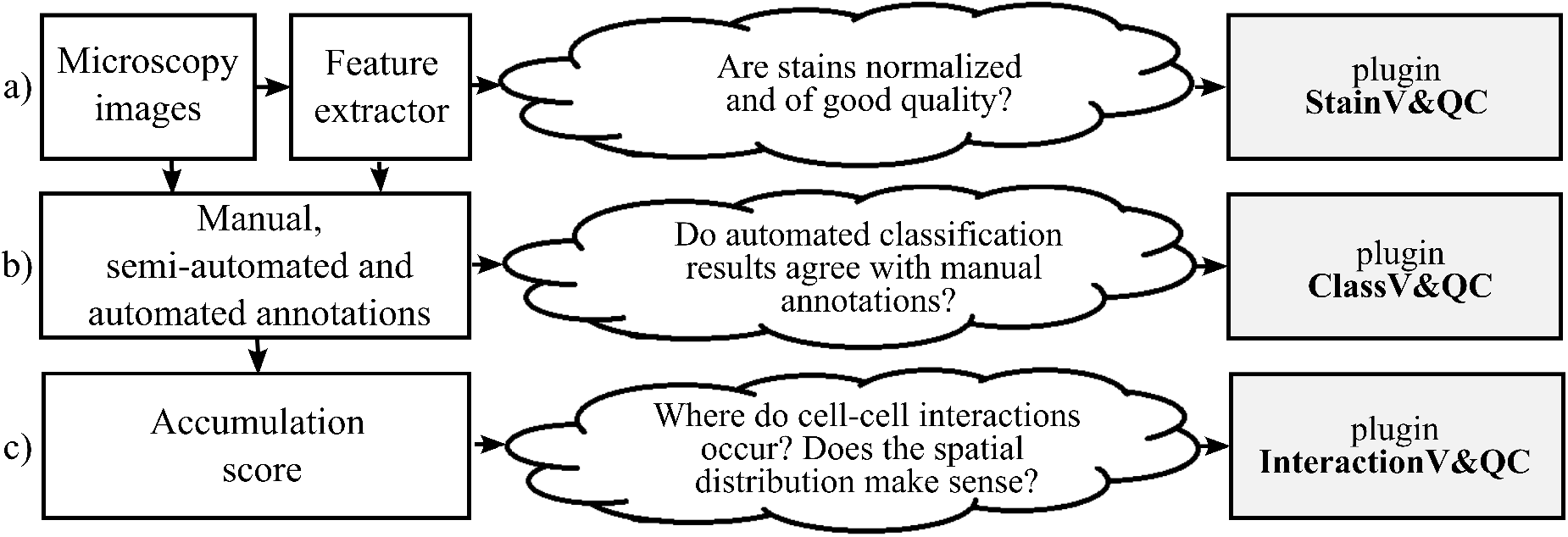
Diagram of the workflow. The left column describes the steps of the analysis, while the center column describes some of the associated quality control questions, and the right column lists the associated visualization and quality control plugins: a) - Plugin for visualization comparison and quality control of cell staining. b) - Plugin for visualization comparison and quality control of cell classification. c) - Plugin for visualization and quality control of cell-cell interactions

### 2.2 Plugins

#### 2.2.1 Visualization comparison and quality control of cell staining (StainV&QC)

Our plugin for visualization comparison and quality control of cell staining, StainV&QC, can visualize a feature space or compare several feature spaces of different samples. With feature space, we refer to the space spanned by all the different measurements extracted from individual cells, such as mean intensity per cell and image channel, but also more advanced features such as measurements of texture and shape. Figure 2 shows the workflow of required steps in order to use the StainV&QC plugin. The first step is to segment individual cells from the multiplexed microscopy data. The second step is to use a feature extractor on segmented cells to extract features. Then either visualize two features at the same time in the plugin or use a dimensionality reduction technique, such as UMAP [25], to visualize all the features but in a lower dimension. It is also possible to visualize several samples at the same time to investigate if their feature spaces match or if some pre-processing steps are necessary to add before further steps. Hypothetically, feature spaces of the same tissue type should approximately match, assuming that at least some portion of the cells in the tissue should be of the same type, and therefore have similar feature spaces. Shifts in feature spaces are typically due to variations in staining and can be corrected by normalization [26]. Such correction is vital for downstream cell classification.

**Figure 2:**
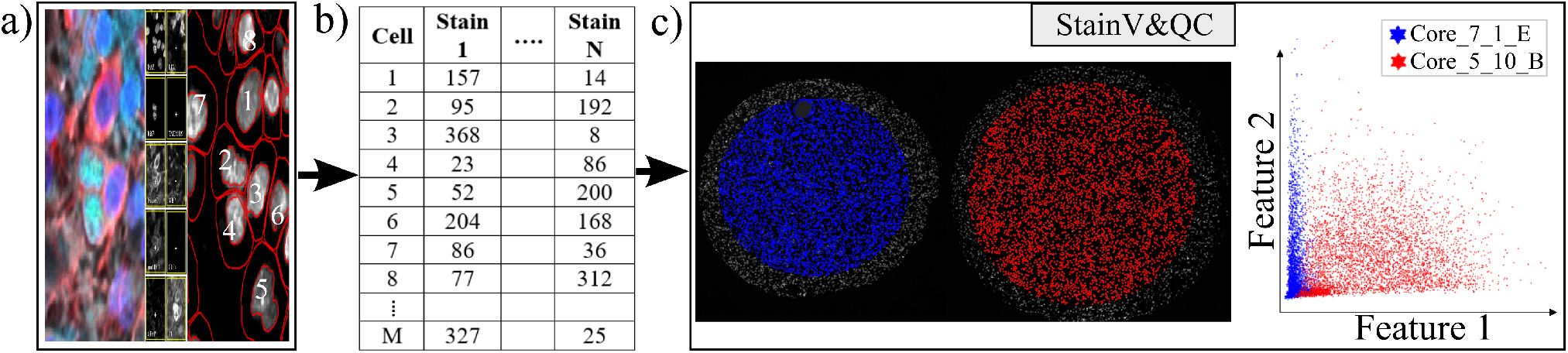
Workflow before using plugin StainV&QC. a) - Multiplexed microscopy data with segmented cells, b) - Table of extracted features from all the segmented cells, c) - Plugin StainV&QC

The main screen of the StainV&QC plugin can be seen in subfigure 2 - c). The left side shows a microscopy image of two tissue samples overlaid by cell markers with different colors per sample, this element is called the Spatial viewport. The right side displays the feature space, where the user can interactively select markers and instantaneously see the corresponding markers in the Spatial viewport. The color of the markers in the feature space is the same as the color of the markers in the Spatial viewport. A detailed description of the plugin settings can be found in the Supplementary material - Figure S1.1. This plugin can be found at https://tissuumaps.github.io/TissUUmaps/plugins/.

StainV&QC can be very useful to identify upstream analysis challenges such as weak staining, high background intensity, over-staining, nonspecific staining, nonuniform illumination in imaging, or low/high contrast.

#### 2.2.2 Visualization comparison and quality control of cell classification (ClassV&QC)

The plugin for visualization, comparison and quality control of cell classification, ClassV&QC, can visualize and set side-by-side results of various techniques for cell classification. Figure 3 shows that the workflow requires that cell classification has first been applied to the cells represented by the feature space. Cell identification (segmentation) followed by cell classification can be done by several approaches, such as traditional manual annotations by an expert, or by semi-automated or automated categorization of the cells by specific tools (e.g. CellProfiler [19, 20] or QuPath [22]). Subsequently, the results are visualized in the ClassV&QC plugin, and different approaches for detection and classification (e.g. manual vs automated or two different automated methods) can be compared and evaluated also in relation to original input data, shown as a mosaic of cut-outs cropped from a region around a selected cell across the channels of the raw data.

**Figure 3:**
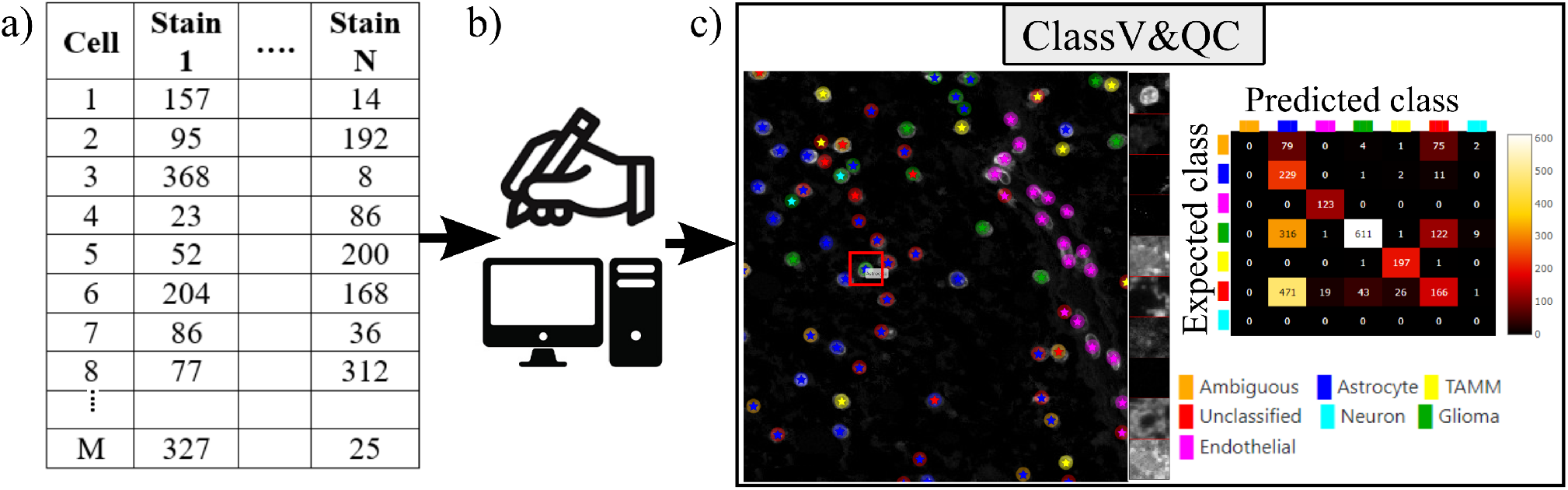
Workflow for using plugin ClassV&QC. a) - Features extracted from the microscopy images are input to manual or automated classification step, b) - Manual, semiautomated, or automated cell classification, c) - Plugin ClassV&QC containing interactive confusion matrix where the user can click on the elements of the matrix and only cells counted in that matrix element are displayed on the Spatial viewport

The main screen of the plugin can be seen in figure 3 - c). The left side shows the Spatial viewport. When comparing two approaches, the result of each approach is presented as a different shape and size of markers. The first approach has bigger circular markers and the second approach has smaller star markers on top of the bigger circular markers. Hence, differences of classification between approaches can be detected easily. The right side of the Spatial viewport shows image patches cropped around a selected cell (red square in Spatial viewport) in all the stain channels of the dataset. This tool can help to investigate potential staining issues associated with cells that are assigned the wrong class. The right side of figure 3 - c) shows an interactive confusion matrix when comparing two cell classification approaches. A confusion matrix is a way to compare the result of two classification results, typically manually or semi-manually annotated cells (expected class), and the results of a fully automated classification method (predicted class). It is also possible to compare the performance of two different automated classification methods. Each row of the matrix represents the elements in an expected class while each column represents the elements in a predicted class. The user can click on the elements of the confusion matrix and only cells counted in that matrix element are displayed on the Spatial viewport. This function requires that the two approaches that are compared have the same cell segmentation/identification as input so that the order of the cells matches. Non-matching cell IDs only enable visualization of cell type distributions and patches of microscopy data. A detailed description of the plugin settings can be found in the Supplementary material - Figure S2.1. This plugin can be found at https://tissuumaps.github.io/TissUUmaps/plugins/.

When using the patches to visually evaluate the quality of the classification, we expect that the objects in the original image data always correspond to the true biological structures which were meant to be imaged. For example, if the stain is supposed to bind to nuclei, we expect to see exclusively nuclei in the final image. The visualization of the patches of the microscopy data then points out false negative classification or wrong classification.

#### 2.2.3 Visualization and quality control of cell-cell interactions (InteractionV&QC)

Cell-cell interaction can be defined as two (or more) cell types with a certain distance to each other that appear with a higher frequency than what would be expected by random distribution. A spatial distribution pattern between two cell types that is statistically significant can indicate involvement in some kind of interaction. For example, immune cells appearing non-randomly close to tumor cells may indicate some kind of interaction. The amount of interaction, and its significance as compared to interaction by chance, can be quantified by several approaches as summarized in the review [16].

Apart from the visualization and quality control plugins, we also present an approach to quantify interactions, using a neighborhood enrichment test (NET) that compares observed cell neighborhoods to randomized patterns. A previous approach to analyzing neighborhood enrichment was presented by Palla et al in SquidPy [23]. The NET presented here automatically compensates for tissue structure variation and can identify cell types that are distributed non-randomly in relation to one another, independent of the intrinsic tissue patterns. A cell neighborhood is defined as all cells within a distance *k* defined by the user. The same distance *k* is used across the whole tissue, independent of cell density. The NET score compares the observed neighborhood relationship between two cell types *A* and *B*, to what could be observed if one of the cell types was randomly distributed. First, the average number of cells of type *B* in the local neighborhood of cell type *A* (defined by the distance *k*) is measured. This gives *n*_*AB*_. Next, the cells of type *B* are randomized to the positions of all other cell types (excluding *A*), while all the cells of type *A* are kept in their original positions. The randomization process is repeated many times (specified by the user, e.g. 1000 times), and for each randomization, the average number of cells of type *B* in the local neighborhood of cell type *A* is measured. Finally, the mean (*μ*_*AB*_) and the standard deviation (*σ*_*AB*_) of cell count averages over all randomizations is calculated, and the NET score is defined as

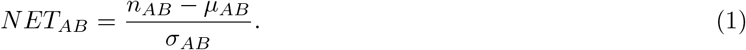

A NET score close to zero indicates that cell types A and B are randomly distributed to one another, while a positive score means that they are attracted to one another, and a negative score indicates that there is a repulsion between the cell types. The output is saved as a csv-file. The code for running the NET analysis can be found as a Jupyter notebook at https://github.com/BIIFSweden/accScore.

Our plugin for visualization and quality control of cell-cell interactions, InteractionV&QC, serves as a tool to visualize and understand non-random tissue patterns and spatial cell-cell interactions, such as those quantified by NET. As presented in figure 4, firstly, cells in microscopy images are classified and once the classification is regarded as valid (e.g. using the ClassV&QC plugin), accumulation scores are calculated by spatial statistic tools, such as NET, Squidpy [23], or histoCAT [24]. The resulting matrix has to be saved as a .csv file for uploading to the InteractionV&QC plugin.

**Figure 4:**
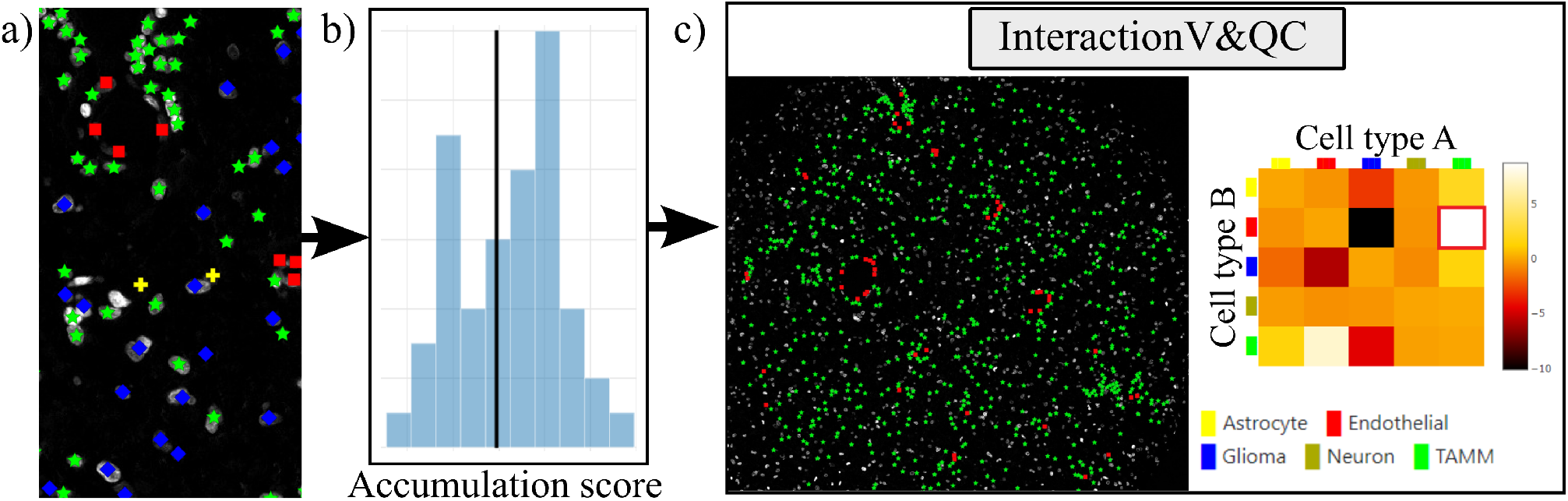
Workflow for using the InteractionV&QC plugin. a) - Cell classification results, b) - Accumulation scores (as quantified using NET), blue bars represent the distribution of the randomized counts of connections and the black line represent the actual count of connections, c) - Plugin InteractionV&QC containing interactive matrix where the user can click on the elements of the matrix and only those two corresponding cell types are displayed on the Spatial viewport

The main screen of the plugin can be seen in figure 4 - c). The left side shows the Spatial viewport and the right side represents a visualization of the neighborhood enrichment test NET. The axes of the matrix are colored based on different cell types as can be seen in the legend, and the color code is the same as the corresponding markers in the Spatial viewport. This matrix is interactive and the user can click on the elements of the matrix and only those two corresponding cell types are displayed on the Spatial viewport. If the interaction between two cell types is quantified as significant, this tool helps to visually explore the interactions and build an understanding of why the interaction may be significant. A detailed description of the plugin settings can be found in the Supplementary material - Figure S3.1. This plugin can be found at https://tissuumaps.github.io/TissUUmaps/plugins/.

InteractionV&QC plugin helps the user to visualize possible non-random interaction between two cell types in the space. This can be done for several different methods of calculating the NET score and the user can visually access which methods’ results make more sense from the spatial point of view.

## 3 Experimental Validation and Results

In this section, we present the application results of the plugins on real-life datasets. We show the plugins’ versatility and usefulness when investigating multiplexed microscopy images.

### 3.1 Results of StainV&QC plugin

For testing the utility of the StainV&QC plugin, we used a dataset from a tissue microarray (TMA) of tumor cores with a diameter of 1.2 mm constructed at the Human Protein Atlas, Department of Immunology, Genetics and Pathology, Uppsala University, Sweden, previously presented in [27]. The TMA underwent octaplex immunofluorescence staining using a 7-plex Opal kit and additional Opal 480 and Opal 780 reagent kits (Akoya Biosciences, Marlborough, US) and DAPI counterstain of cell nuclei. Images were obtained after scanning the TMA through Vectra Polaris (Akoya Biosciences) at 20× magnification. For this experiment, we selected two TMA cores and used their corresponding DAPI images as input for QuPath to segment all cell nuclei. We also created an approximate cell segmentation by dilation of these cell nuclei. These areas were used to extract features per cell by QuPath. The features are basic statistics of intensities (mean, min, max, and SD). The description of an example file can be found in the Supplementary material - Figure S1.2.

Before proceeding to cell classification, we loaded the data to the StainV&QC plugin. As can be seen in figure 5 - middle, the feature spaces are distinct from each other which means very strong differences in the image intensities between the two cores. These differences may be due to variations in tissue fixation, leading to variations in antibody binding properties. Next, we applied normalization of extracted features through Winsorization [28], defined as

**Figure 5:**
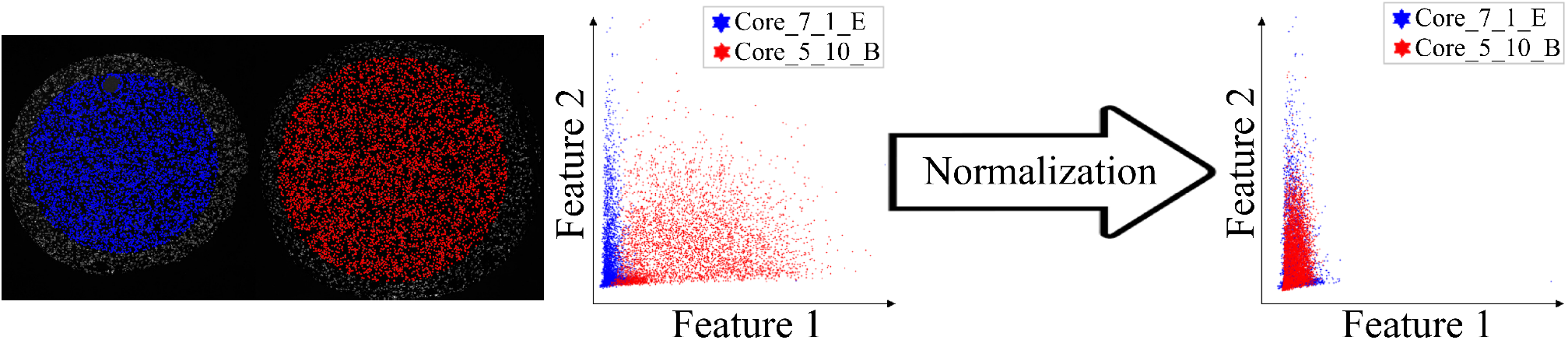
Main screen of StainV&QC plugin, comparing two cores with corresponding data in feature space before and after normalization. Colors represent individual tissue cores

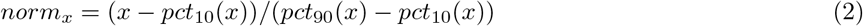

Here, *pct*_10_ and *pct*_90_ are the 10th and 90th percentiles of the corresponding feature measurements *x*. The normalized data was loaded to the StainV&QC plugin and we got aligned feature spaces as can be seen in figure 5 - right. Using the StainV&QC plugin, we can quickly confirm that the aim of the normalization step, to make the feature distribution of the two cores more equal, has been reached. This is important for the following steps involving automated cell classification.

### 3.2 Results of ClassV&QC plugin

For showcasing the ClassV&QC plugin, we used the same dataset presented above, and the ClassV&QC plugin was used to compare two different cell classification results. The first classification result (referred to as *Expected class*) consists of manual annotations by an expert and is described in the Materials section of [27], and the second result (referred to as *Predicted class*) is the result of a fully automated classification result based on a fully convolutional neural network (FCNN) model [27]. Both classification approaches start from the same cell segmentation/identification step as presented above. The description of the example files can be found in the Supplementary material - Figure S2.2.

Figure 6 - left shows the classification results of both approaches, circles represent the results of manual annotations colored by cell type and stars represent the results of FCNN classification colored by cell type. This visualization helps to investigate where selected methods match and mismatch from the spatial point of view. In cases where the methods mismatch, the ClassV&QC plugin makes it easy to investigate which result is correct by clicking on the cell marker to visualize patches from all the image channels from the raw microscopy data. These patches are shown on the right side of the Spatial viewport. For instance, the selected cell was manually classified as a Glioma cell, corresponding to high-intensity values in the Opal650 image channel marking mutIDH1. However, the FCNN algorithm classified the same cell as Astrocyte, meaning there should be higher intensity in the Opal780 channel marking GFAP and no signal from mutIDH1. Then the user can use the ClassV&QC plugin to visually evaluate which method has a more reliable classification result. The right side of figure 6 shows an interactive confusion matrix that visualizes a comparison of the classification results by the two approaches. The user can interactively click on the matrix’s elements to view only specific disagreements or only specific agreements between the cells on the Spatial viewport. This can be used, for example, to investigate if certain disagreements are significant only for a particular area of the tissue.

**Figure 6:**
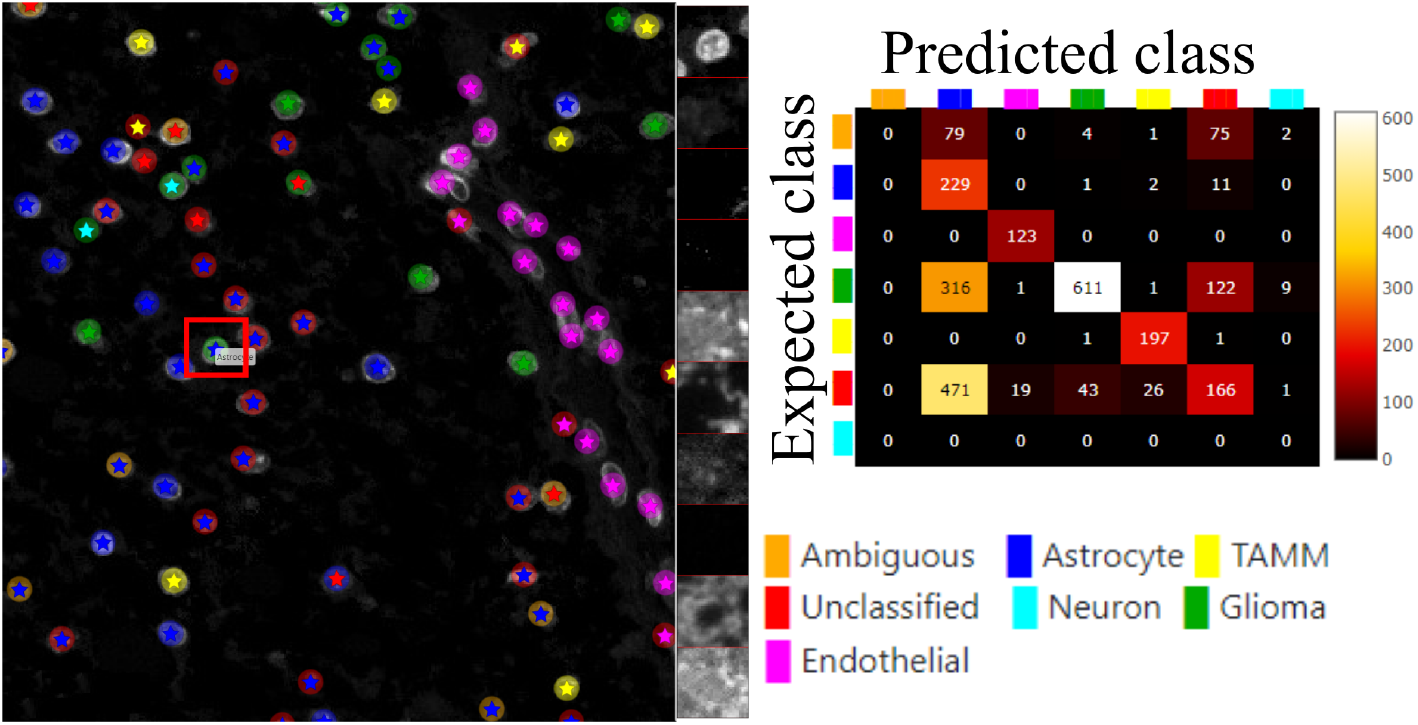
Main screen of the ClassV&QC plugin, comparing two classification techniques in the Spatial viewport with a corresponding confusion matrix. Circles represent the results of manual annotations colored by cell type and stars stand for FCNN classification colored by cell type

In order to further verify the value of the ClassV&QC plugin, we also tested it on fluorescence microscopy images from the open dataset originally provided by Ilya Ravkin and made publicly available via the Broad Bioimage Benchmark Collection [29]. The images are from a drug screening experiment, where human U2OS cells were grown in a 96-well plate with varying doses of two drugs. As the drug dose increases, a protein tagged with the green fluorescent protein GFP is translocated from the cytoplasm to the nucleus, and thus the amount of GFP expressed in the nuclei increases and GFP expressed in the cytoplasm decreases. The goal of the analysis is to quantify this translocation of GFP, or more specifically, to measure the fraction of cells in an image that have nuclear or cytoplasmic GFP expression. In the evaluation of the ClassV&QC plugin, we compared results achieved by two different, fully automated classification methods; SimSearch [30] and CellProfiler [19, 20]. CellProfiler classifies cells by first identifying individual cell nuclei, and then extracting measurements such as staining intensities from the surrounding area to assign cell classes. SimSearch is based on deep learning, and searches for image patches that fit a pattern learned from examples of cells of different classes. This means that there may not be a 1:1 match between cell IDs in SimSearch and CellProfiler. More detailed instructions on how to use the plugin, as well as illustrations of example files can be found in the Supplementary material - Figure S2.3.

Figure 7 shows the results of these two classification methods; discs represent CellProfiler results colored by the cell category and stars stand for SimSearch results colored by its proposed cell class category. In this case, there are three cell categories: GFP in the nucleus (purple), GFP in the cytoplasm (orange), and no GFP (green). Cropped patches from each staining image, specifically the distribution of GFP (top) and a nuclear stain (bottom) are displayed in the right corner of the viewport. Using the ClassV&QC plugin, it is easy to see that the selected cell has high GFP intensities localized to the cytoplasm, so it has been correctly classified as having GFP in the cytoplasm by SimSearch (orange star), while CellProfiler incorrectly classified it as having GFP in the nucleus (purple disc).

**Figure 7:**
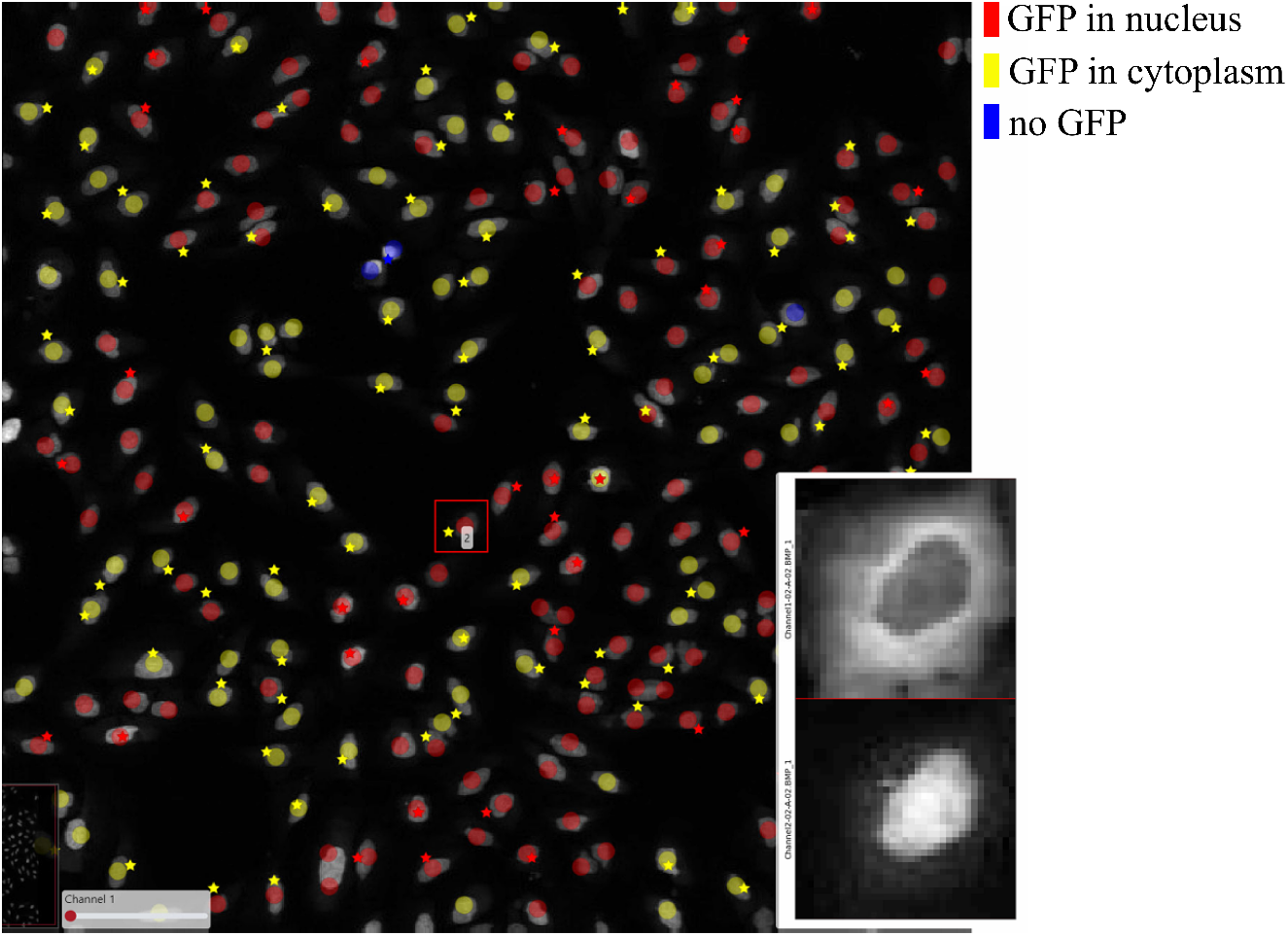
Main screen of ClassV&QC plugin, comparing two classification techniques applied to human U2OS cell cultures. Discs represent CellProfiler results colored by the cell category and stars stand for SimSearch results colored by the cell category

### 3.3 Results of InteractionV&QC plugin

Finally, we evaluated the usefulness of the InteractionV&QC plugin. For this experiment, we once again used the multiplexed immunofluorescence dataset presented above. We selected one TMA core containing all five possible cell types and then we used the NET score Jupyter Notebook described above to calculate the neighborhood enrichment between all combinations of the cell type pairs and saved it as a .csv file. The description of the example file can be found in the Supplementary material - Figure S3.2. Subsequently, the .csv file was loaded into the InteractionV&QC plugin to visualize the matrix, as can be seen in figure 8 - a). The user can interactively click on any element of the matrix and it displays two corresponding cell types in the Spatial viewport. In figure 8 we selected the test results between Glioma (blue) and tumor-associated macrophages/microglia (TAMM) (green) cells since the test results show significant interaction (positive NET score). The InteractionV&QC makes it possible to instantaneously observe the corresponding cell types in the Spatial viewport, highlighting the spatial location of the interaction of these two cell types.

**Figure 8:**
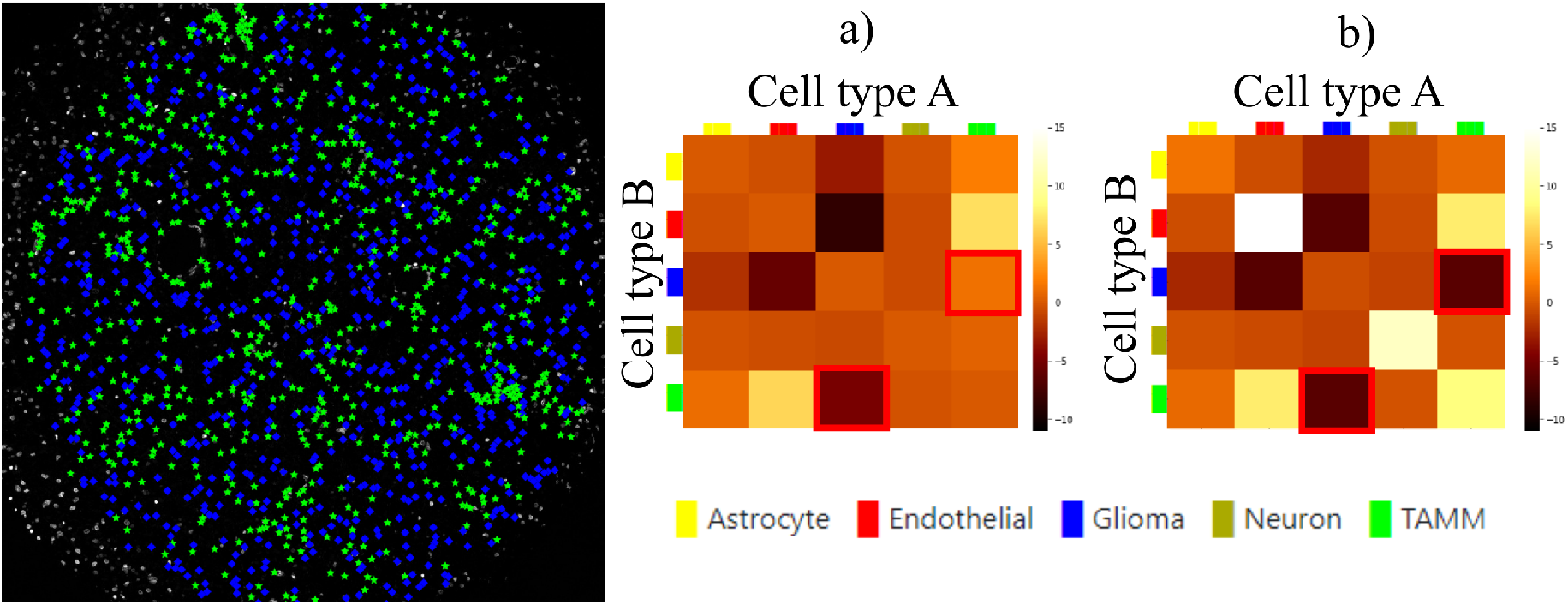
The main screen of the InteractionV&QC plugin. a) - neighborhood enrichment calculated by NET, b) - neighborhood enrichment calculated by Squidpy

To illustrate the utility of the tool, we compare neighborhood enrichments calculated by the NET score and Squidpy as shown in figure 8 - b). The difference is that when using our NET method for calculating the accumulation score, we keep the locations and labels of one cell type and shuffle all the other ones. In the implementation of Squidpy, all the cell locations are shuffled, which is faster. However, we would like to argue that in the case of the NET method presented here, we automatically compensate for the presence of tissue structures, such as vessels, or tissue samples section containing more than one tissue type. By keeping the locations of one cell type which is appearing only in one part of the tissue (one tissue type), we do not create false indications of repulsion or interaction when randomizing the remaining cell types in the computation of the accumulation score. If we look at the highlighted matrix elements in figure 8 - a) and b), we can see that our NET score is asymmetrical indicating that the green TAMM cells are repulsed in relation to the Glioma cells (the TAMM cell are clustered and the NET score is negative), while the Glioma cells are randomly distributed in relation to the TAMM cells (NET score close to zero). The Squidpy implementation does not pick up these differences.

## 4 Conclusion

It is important to note that it is not possible to extract quantitative results of quality using these plugins. The plugins serve as a means of visual quality control, and not as a processing step. Quantitative quality control relies on a set of problem-specific definitions of what is considered ‘good enough’. For example, in figure 7, one could argue that the result is ‘good enough’ if the cell count of the two methods is within a certain error range, or one could argue that the distance between the location of each cell of a given class must be within a certain minimum distance of cells of the same class as presented by the other method. And in the end, one may want to compare to visual annotations, which are very expensive to produce. A fast and efficient tool, such as TissUUmaps, for visually presenting results, can give the user quick insights into quality, without tedious annotations. Visual assessment can also be a valuable tool in designing metrics for quantitative evaluation.

To conclude, the visualization and quality control tools presented here have the potential to function as an important bridge between visual/manual assessment and fully automated approaches to quantitatively extract information from large-scale multiplex microscopy experiments. Since truly validated ground truth is lacking in this type of assay, tools for interactive quality control are necessary for methods comparison, optimization, and validation. The plugins presented here are part of the free and open-source TissUUmaps 3 project, designed to scale to very large datasets. Thanks to access via a web browser, it enables easy sharing of the plugins and data in multi-disciplinary projects across different labs, without having to transfer data. We believe these visualization and quality control plugins will forward the field through efficient optimization and building of trust in automated analysis, enabling large-scale studies advancing our understanding of medicine and biology.

## Acknowledgements

We are grateful for sharing access to data and technical assistance of Anja Smits, Thomas Olsson Bontell and Asgeir Jakola.

## Data Availability Statement

The code can be found at https://tissuumaps.github.io/TissUUmaps/plugins/.

## Ethical Standards

The research meets all ethical guidelines, including adherence to the legal requirements of the study country.

## Author Contributions

Conceptualization: A.B., C.W.; Data curation, L.W., A.O.; Funding acquisition, C.W., A.O.; Investigation, A.B., E.C., A.A., C.W.; Methodology, A.B., C.A., A.A., E.C., A.K., C.W.; Project administration, C.W.; Resources, C.W., A.O.; Software, A.B., C.A., E.C.; Supervision, C.W., A.O.; Visualization, A.B., C.A.; Writing—original draft, A.B.; Writing—review & editing, A.B., C.A., A.A., E.C., A.K., L.W., A.O., C.W. All authors approved the final submitted draft.

## Supplementary Material

The supplementary material provides a detailed description of the settings of the plugins.

