## Supplementary material for "Visualization & Quality Control Tools for Large-scale Multiplex Tissue Analysis in TissUUmaps 3": Suplementary material

The plugins presented in the paper will automatically show up in the 'Plugins' menu of TissUUmaps, which can be downloaded from https://tissuumaps.github.io/download/

Check the boxes in front of the plugins of interest, click 'Install', and re-start TissUUmaps. Next, parameters are set as described below.

#### Plugin StainV&QC:

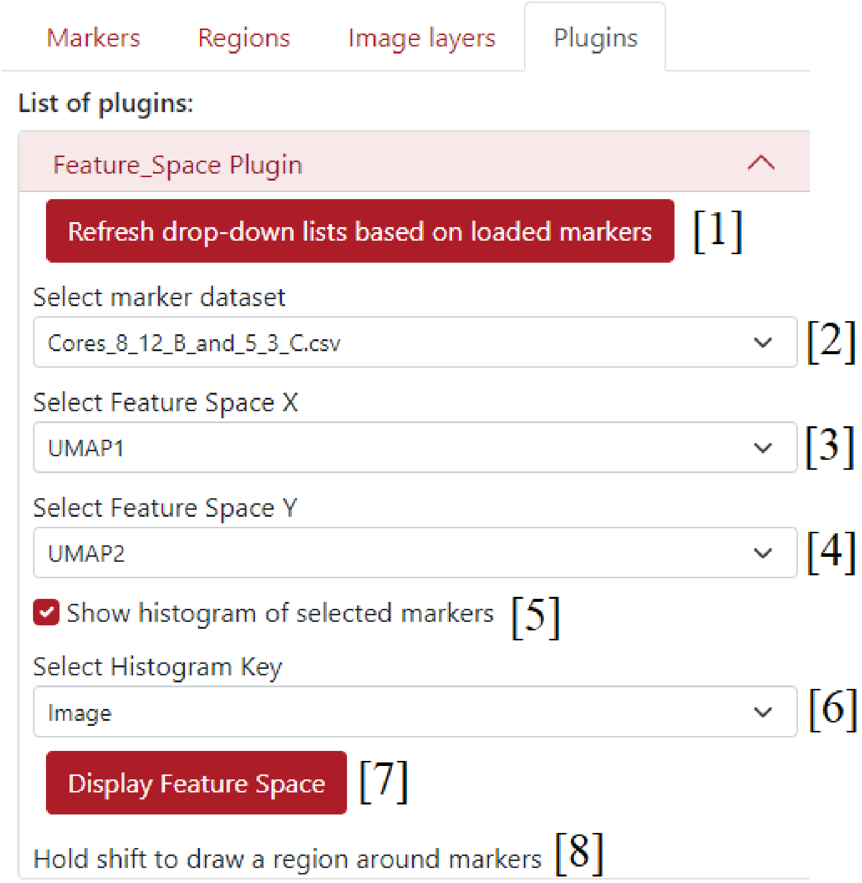


Supplementary figure S1.1: Settings of StainV&QC plugin; [1] - button *Refresh drop-down lists based on loaded markers* - to update the pop-up menus to all the loaded marker files, [2] - pop-up menu *Select marker dataset* - to select which dataset will be used to choose the feature space coordinates, [3] - pop-up menu *Select Feature Space X* - to select the feature spaces X column from previously selected dataset in [2], [4] - pop-up menu *Select Feature Space Y* - to select the feature spaces Y column from previously selected dataset in [2], [5] - checkbox *Show histogram of selected markers* - to give an option to show or not the histogram of selected markers, [6] - pop-up menu *Select Histogram Key* - to select a key column used to calculate the histogram, [7] - button *Display Feature Space* - to use the selected feature spaces X and Y as Cartesian coordinates to display the feature space as in Spatial viewport, [8] - info *Hold shift to draw a region around markers* - info to draw regions in the feature space.

| 1 | Centroid X | Centroid Y | Nucleus: Area | Nucleus: Perimeter | Nucleus: Circularity | Nucleus: Max caliper | Nucleus: Min caliper | Nucleus: Eccentricity |
|---|---|---|---|---|---|---|---|---|
| 2 | 1058.4 | 333.15 | 41.5 | 25.7055 | 0.7892 | 9.4683 | 5.6842 | 0.819 |
| 3 | 951.9 | 331.74 | 41 | 26.0766 | 0.7577 | 9.7378 | 5.6042 | 0.8369 |
| 4 | 1094.2 | 332.36 | 9.5 | 13.9851 | 0.6104 | 6.0092 | 2.3496 | 0.9257 |
| 5 | 1115.3 | 336.93 | 44.25 | 25.9065 | 0.8285 | 8.626 | 7.5329 | 0.4234 |
| 6 | 968.12 | 337.85 | 31.5 | 20.726 | 0.9215 | 7.5258 | 5.604 | 0.6775 |
| 7 | 1142.4 | 339.25 | 14.25 | 15.7015 | 0.7263 | 6.492 | 2.9525 | 0.8816 |
| 8 | 893.32 | 341.36 | 31.25 | 21.1487 | 0.878 | 7.488 | 5.2116 | 0.6986 |
| 9 | 948.5 | 340.91 | 17 | 15.1324 | 0.9329 | 5.5187 | 3.5559 | 0.7584 |
| 10 | 862.83 | 346.65 | 16 | 16.1047 | 0.7752 | 6.1609 | 3.2184 | 0.8688 |
| 11 | 1041 | 349.07 | 34.5 | 23.9652 | 0.7549 | 9.7845 | 4.7712 | 0.8754 |
| 12 | 889.69 | 350.94 | 38.75 | 23.6156 | 0.8731 | 8.4548 | 6.1036 | 0.7463 |
| 13 | 1148.2 | 350.8 | 40.5 | 23.3984 | 0.9296 | 8.832 | 5.9751 | 0.7358 |
| 14 | 929.22 | 353.6 | 55.25 | 30.2706 | 0.7577 | 12.5895 | 6.3378 | 0.8748 |
| 15 | 856.59 | 354.78 | 65 | 32.5622 | 0.7704 | 10.7927 | 8.9944 | 0.4843 |
| 16 | 904.8 | 352.83 | 47.25 | 24.7836 | 0.9667 | 8.25 | 7.4477 | 0.3793 |
| 17 | 811.25 | 354.36 | 24 | 19.3194 | 0.808 | 7.7385 | 4.5499 | 0.7997 |

Supplementary figure S1.2: Example of an input dataset for StainV&QC plugin. This example follows the standard csv-file output from software like QuPath (1) and CellProfiler (2). The first two columns contain X and Y coordinates of each object (in this case nuclei), and the remaining columns contain features extracted from the original microscopy image, such as nucleus perimeter or nucleus eccentricity. These features can be selected in the plugin's settings *Select Feature Space X* and *Select Feature Space Y* as shown in the Figure S1.1. File rows represent markers (data observations), in this case individual cells.

### Plugin ClassV&QC:

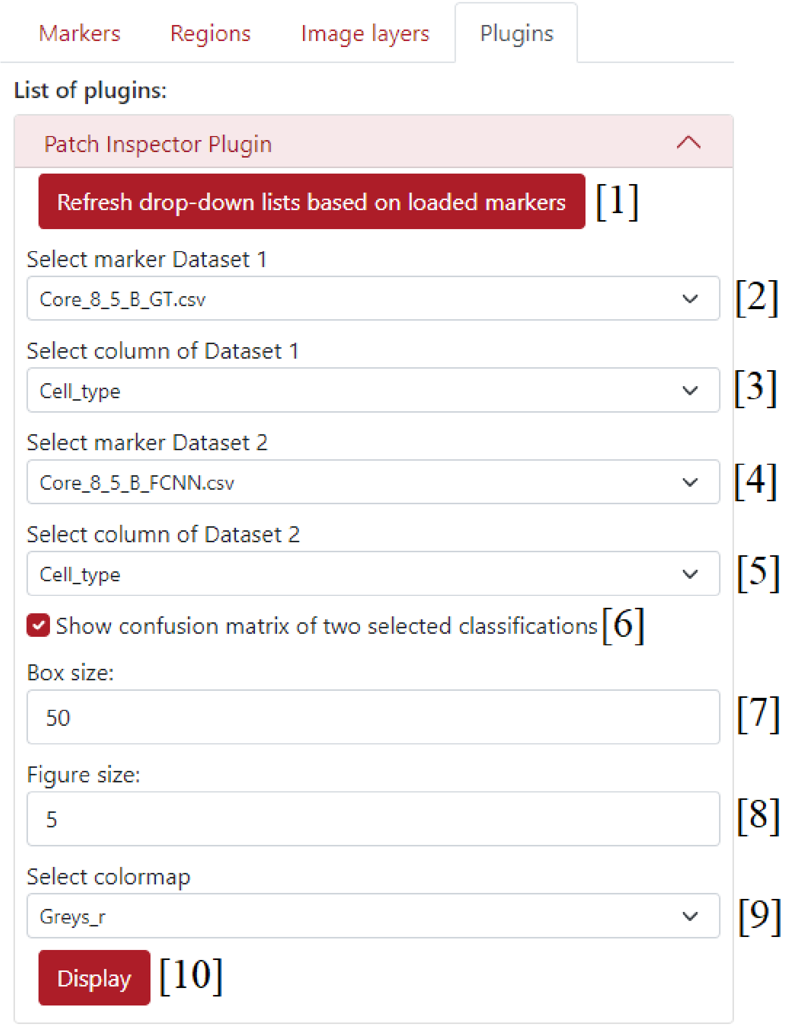


Supplementary figure S2.1: Settings of ClassV&QC plugin; [1] - button *Refresh drop-down lists based on loaded markers* - to update the pop-up menus to all the loaded marker files, [2] - pop-up menu *Select marker Dataset 1* - to select the first marker datasets for the comparison of different techniques in spatial viewport, [3] - pop-up menu *Select column of Dataset 1* - to select the column of the Dataset 1 for the comparison of different techniques in spatial viewport, [4] - pop-up menu *Select marker Dataset 2* - to select the second marker datasets for the comparison of different techniques in spatial viewport, [5] - pop-up menu *Select column of Dataset 2* - to select the column of the Dataset 2 for the comparison of different techniques in spatial viewport, [6] - checkbox *Show confusion matrix* - if it is checked, the confusion matrix is calculated and displayed underneath the settings, [7] - *Box size* - to select the width and height in pixels of the image patch cropped from the microscopy image represented by a red square in the Spatial viewport, [8] - *Figure size* - to determine the width of the displayed column of the patches, [9] - pop-up menu *Select colormap* - to select the colormap for the visualization of image patches, [10] - button *Display* - to use selected datasets and display them in the Spatial viewport and optional confusion matrix.

a)

| 1 | X | Y | Cell_type |
|---|---|---|---|
| 2 | 1874.231 | 168 | Unclassified |
| 3 | 1848.231 | 165.0769 | Unclassified |
| 4 | 1953.286 | 167.3571 | Ambiguous |
| 5 | 1968.389 | 176.6667 | Glioma |
| 6 | 1839.4 | 178.9 | Glioma |
| 7 | 1710 | 175.4545 | Ambiguous |
| 8 | 1985.875 | 183 | Unclassified |
| 9 | 2070.867 | 185.6 | Glioma |
| 10 | 2090.231 | 187 | Glioma |
| 11 | 2131.444 | 185.1111 | Endothelial |
| 12 | 2119.333 | 187.2222 | Endothelial |
| 13 | 2141.533 | 193.2667 | Endothelial |

b)

| 1 | X | Y | FCNN |
|---|---|---|---|
| 2 | 1874.231 | 168 | TAMM |
| 3 | 1848.231 | 165.0769 | Endothelial |
| 4 | 1953.286 | 167.3571 | Astrocyte |
| 5 | 1968.389 | 176.6667 | Astrocyte |
| 6 | 1839.4 | 178.9 | Glioma |
| 7 | 1710 | 175.4545 | Astrocyte |
| 8 | 1985.875 | 183 | Unclassified |
| 9 | 2070.867 | 185.6 | Glioma |
| 10 | 2090.231 | 187 | Unclassified |
| 11 | 2131.444 | 185.1111 | Endothelial |
| 12 | 2119.333 | 187.2222 | Endothelial |
| 13 | 2141.533 | 193.2667 | Endothelial |

Supplementary figure S2.2: Example of input datasets for ClassV&QC plugin; a) – shows an example of the .csv file containing ground truth data, this file needs to contain columns of X and Y coordinates and an identity column (Cell_type). b) – shows an example of the .csv file containing classification results from the fully connected neural network (FCNN), this file needs to contain columns of X and Y coordinates and an identity column (FCNN). You can see that the X and Y coordinates of individual observations (rows) match over example datasets a) and b). This allows the plugin to use the feature of confusion matrix visualization because the cell IDs match.

a)

| 1 | Cells_Location_Center_X | Cells_Location_Center_Y | CellID |
|---|---|---|---|
| 2 | 353.8937947 | 623.9045346 | 2 |
| 3 | 104.2520107 | 626.7587131 | 0 |
| 4 | 439.463094 | 625.0323559 | 0 |
| 5 | 599.1293833 | 622.8222491 | 2 |
| 6 | 305.3108571 | 621.4857143 | 0 |
| 7 | 142.3970588 | 620.8272059 | 2 |
| 8 | 180.0721112 | 614.8792354 | 2 |
| 9 | 484.7188889 | 608.7622222 | 0 |
| 10 | 216.380719 | 609.2287582 | 2 |
| 11 | 76.79185939 | 599.1887142 | 0 |

b)

| 1 | Class ID | X center | Y center |
|---|---|---|---|
| 2 | 0 | 24 | 76 |
| 3 | 0 | 24 | 132 |
| 4 | 0 | 24 | 196 |
| 5 | 0 | 28 | 468 |
| 6 | 0 | 36 | 236 |
| 7 | 0 | 44 | 96 |
| 8 | 0 | 44 | 344 |
| 9 | 0 | 48 | 56 |
| 10 | 0 | 52 | 328 |
| 11 | 0 | 64 | 120 |

Supplementary figure S2.3: Example of input datasets for ClassV&QC plugin; a) – shows an example of the .csv file CellProfiler (2) classification results, this file needs to contain columns of X and Y coordinates and an identity column (CellID). b) – shows an example of the .csv file containing SimSearch (3) classification results, this file needs to contain columns of X and Y coordinates and an identity column (Class ID). You can see that the X and Y coordinates of individual observations (rows) **do not** match over example datasets a) and b). This is the reason why the user can't use the feature of confusion matrix visualization.

Further instructions for how to use this plugin can be found at https://github.com/TissUUmaps/TissUUmaps/blob/master/examples/Instructions%20for%20using%20plugins/2_ClassV%26QC_plugin_in_TissUUmaps.md

### Plugin InteractionV&QC:

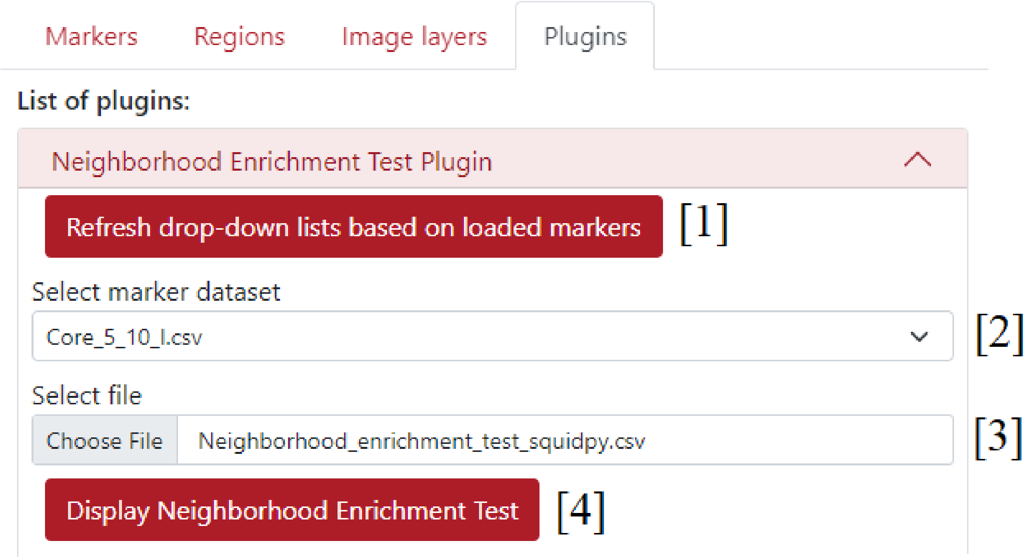


Supplementary figure S3.1: Settings of InteractionV&QC plugin; [1] - button *Refresh drop-down lists based on loaded markers* - to update the pop-up menus to all the loaded marker files, [2] - pop-up menu *Select marker dataset* - to select the marker dataset which will be used for the interactive visualization of the neighborhood enrichment test matrix, [3] - *Select file* - to select a .csv file from the computer which contains a matrix of neighborhood enrichment test results, [4] - button *Display Neighborhood Enrichment Test* - to load the .csv file and displays interactive matrix underneath the settings.

| | A | B | C | D | E | F |
|---|---|---|---|---|---|---|
| 1 | 0 | Astrocyte | Endothelia | Glioma | Neuron | TAMM |
| 2 | Astrocyte | 0.904428 | -0.59821 | -2.25947 | -0.49975 | -0.05219 |
| 3 | Endothelia | 0.575801 | 14.01056 | -8.60607 | -0.85505 | 5.947993 |
| 4 | Glioma | -2.49383 | -11.6682 | 8.220385 | -0.53346 | -7.08637 |
| 5 | Neuron | -0.51427 | -0.83298 | -0.43486 | 7.971367 | -0.5463 |
| 6 | TAMM | 0.724889 | 4.938696 | -6.00065 | -0.20416 | 5.42688 |
| 7 | | | | | | |

Supplementary figure S3.2: Example of input datasets for InteractionV&QC plugin; it contains results of neighborhood enrichment test (NET) in a form of matrix, each matrix element represents proximity score between two cell types listed in the first row and first columns of the file. For example, element *B3* scores 0.575801 in NET between Astrocytes and Endothelial cells.

Further instructions for how to use this plugin can be found at [https://github.com/TissUUmaps/TissUUmaps/blob/master/examples/Instructions%20for%20using%20plugins/3_InteractionV&QC_plugin_in_TissUUmaps.md](https://github.com/TissUUmaps/TissUUmaps/blob/master/examples/Instructions%20for%20using%20plugins/3_InteractionV&QC_plugin_in_TissUUmaps.md)
